# Abstract concept learning in a simple neural network inspired by the insect brain

**DOI:** 10.1101/268375

**Authors:** Alex J. Cope, Eleni Vasilaki, Dorian Minors, Chelsea Sabo, James A.R. Marshall, Andrew B. Barron

## Abstract

The capacity to learn abstract concepts such as ‘sameness’ and ‘difference’ is considered a higher-order cognitive function, typically thought to be dependent on top-down neocortical processing. It is therefore surprising that honey bees apparantly have this capacity. Here we report a model of the structures of the honey bee brain that can learn sameness and difference, as well as a range of complex and simple associative learning tasks. Our model is constrained by the known connections and properties of the mushroom body, including the protocerebral tract, and provides a good fit to the learning rates and performances of real bees in all tasks, including learning sameness and difference. The model proposes a novel mechanism for learning the abstract concepts of ‘sameness’ and ‘difference’ that is compatible with the insect brain, and is not dependent on top-down or executive control processing.

---

Abstract concepts involve the relationships between things. Two simple and classic examples of abstract concepts are ‘sameness’ and ‘difference’. These categorise the relative similarity of things: they are properties of a relationship between objects, but they are independent of, and unrelated to, the features of the objects themselves. The capacity to identify and act on abstract relationships is a higher-order cognitive capacity, and one that is considered critical for any operation involving equivalence or general quantitative comparison (Wright and Katz, 2007; Piaget and Inhelder, 1969; Daehler and Greco, 1985; Avarguès-Weber and Giurfa, 2013). The capacity to recognise abstract concepts such as sameness has even been considered to form the “very keel and backbone of our thinking” (James, 1890). Several non-verbal animals have been shown to be able to recognise ‘sameness’ and ‘difference’ including, notably, the honey bee (Wright, 1997, 1992; Giurfa et al., 2001; D’Amato et al., 1985).

The ability of the honey bee to recognise ‘sameness’ and ‘difference’ is interesting, as the learning of abstract concepts is interpreted as a property of the mammalian neocortex, or of regions of the avian pallium (Diekamp et al., 2002; Wallis et al., 2001; Miller et al., 2003) and to be a form of top-down executive modulation of lower-order learning mechanisms (Avarguès-Weber and Giurfa, 2013; Miller et al., 2003). This interpretation has been reinforced by the finding that activity of neurons in the prefrontal cortex of rhesus monkeys (*Macaca mulatta*) correlates with success in recognising sameness in tasks (Wallis et al., 2001; Miller et al., 2003). The honey bee, however, has nothing like a prefrontal cortex in its much smaller brain.

In this paper we use a modelling approach to explore how an animal like a honey bee might be able to solve an abstract concept learning task. To consider this issue we must outline in more detail how learning of sameness and difference has been demonstrated in honey bees, and originally in other animals.

A family of ‘match-to-sample’ tasks has been developed to evaluate sameness and difference learning in non-verbal animals. In these tasks animals are shown a sample stimulus followed, after a delay, by two stimuli: one that matches the sample and one that does not. Sometimes delays of varying duration have been imposed between the presentation of the sample and matching stimuli to test duration of the ‘working memory’ required to perform the task (Wright and Katz, 2007; Katz et al., 2007). This working memory concept is likened to a neural scratchpad that can store a short term memory of a fixed number of items, previously seen but no longer present (Baddeley and Hitch, 1974). Tests in which animals are trained to choose matching stimuli are described as Match-to-Sample (MTS) or Delayed-Match-To-Sample (DMTS) tasks, and tests in which animals are trained to choose the non-matching stimulus are Not-Match-To-Sample (NMTS) or Delayed-Not-Match-To-Sample (DNMTS) tasks.

On their own, match-to-sample tasks are not sufficient to show concept learning of sameness or difference. For this it is necessary to show, having been trained to select matching or non-matching stimuli, that the animal can apply the concept of sameness or difference in a new context (Avarguès-Weber and Giurfa, 2013). Typically this is done by training animals with one set of stimuli and testing whether they can perform the task with a new set of stimuli (Wright, 1997, 1992; Giurfa et al., 2001; D’Amato et al., 1985); this is referred to as a transfer test.

In a landmark study Giurfa et al. (2001) showed that honey bees can learn both sameness and difference. They could learn both DMTS and DNMTS tasks and generalise performance in both tasks to tests with new, previously unseen, stimuli (Giurfa et al., 2001). In this study free-flying bees were trained and tested using a Y-maze in which the sample and matching stimuli were experienced sequentially during flight, with the sample at the entrance to the maze and the match stimuli at each of the y-maze arms. Bees could solve and generalise both DMTS and DNMTS tasks when trained with visual stimuli, and could even transfer the concept of sameness learned in an olfactory DMTS task to a visual DMTS task, showing cross-modal transfer of the learned concept of sameness (Giurfa et al., 2001). Bees took 60 trials to learn these tasks (Giurfa et al., 2001); this is much longer than learning a simple olfactory or visual associative learning task, which can be learned by bees in 3 trials (Matsumoto et al., 2012). Their performance in DMTS and DNMTS was not perfect either; the population average for performance in test and transfer tests was around 75%, but they could clearly perform at better than chance levels (Giurfa et al., 2001) in both.

The concept of working memory is crucial for solving a DMTS/DNMTS task, as information about the sample stimulus is no longer available externally to the animal when choosing between the match stimuli. If there is no neural information that can identify the match then the task cannot be solved. We therefore must identify in the honeybee a candidate for providing this information in order to produce a model that can solve the task.

A previous model by Arena et al. (2013) demonstrates DMTS and DNMTS with transfer, however the model contains many biologically unfounded mechanisms that are solely added for the purpose of solving these tasks, and the outcome of these additions disagrees with neurophysiological, and behavioural evidence. We instead take an approach of constraining our model strongly to established neurophysiology and neuronanatomy, and demonstrating behaviour that matches that of real bees. We will compare this model to the model presented here further in the Discussion.

The honey bee brain is structured as discrete regions of neuropil (zones of synaptic contact). These are well described, as are the major tracts connecting them (Strausfeld, 2012). The learning pathways have been particularly intensely studied (*e.g.* Menzel, 2001; Søvik et al., 2015; Giurfa, 2007; Galizia, 2014). The mushroom bodies (*corpora pedunculata*) receive processed olfactory, visual and mechanosensory input (Mobbs, 1982) and are a locus of multimodal associative learning in honey bees (Menzel, 2001). They are essential for reversal and configural learning (Avarguès-Weber and Giurfa, 2013; Boitard et al., 2015; Devaud et al., 2015). Avarguès-Weber and Giurfa (Avarguès-Weber and Giurfa, 2013) have argued the mushroom bodies to be the most likely brain region supporting concept learning, because of their roles in stimulus identification, classification and elemental learning (Galizia, 2014; Bazhenov et al., 2013; Menzel, 2001). Yet it is not clear how mushroom bodies and associated structures might be able to learn abstract concepts that are independent of any of the specific features of learned stimuli and, crucially, how the identity of the sample stimulus could be represented. Solving such a problem requires two computational components. First, a means of storing the identity of the sample stimulus, a form of working memory; second, a mechanism that can learn to use this stored identity to influence the behaviour at the decision point. Below we propose a model of the circuitry of the honey bee mushroom bodies that can perform these computations and is able to solve DMTS and DNMTS tasks.

## Results

### Key model principles: A circuit model inspired by the honey bee mushroom bodies

We explored whether a neural circuit model, inspired and constrained by the known connections of the honey bee mushroom bodies, is capable of learning sameness and difference in a DMTS and DNMTS task (Figure 1). Full details of the models can be found in Methods.

**Figure 1:**
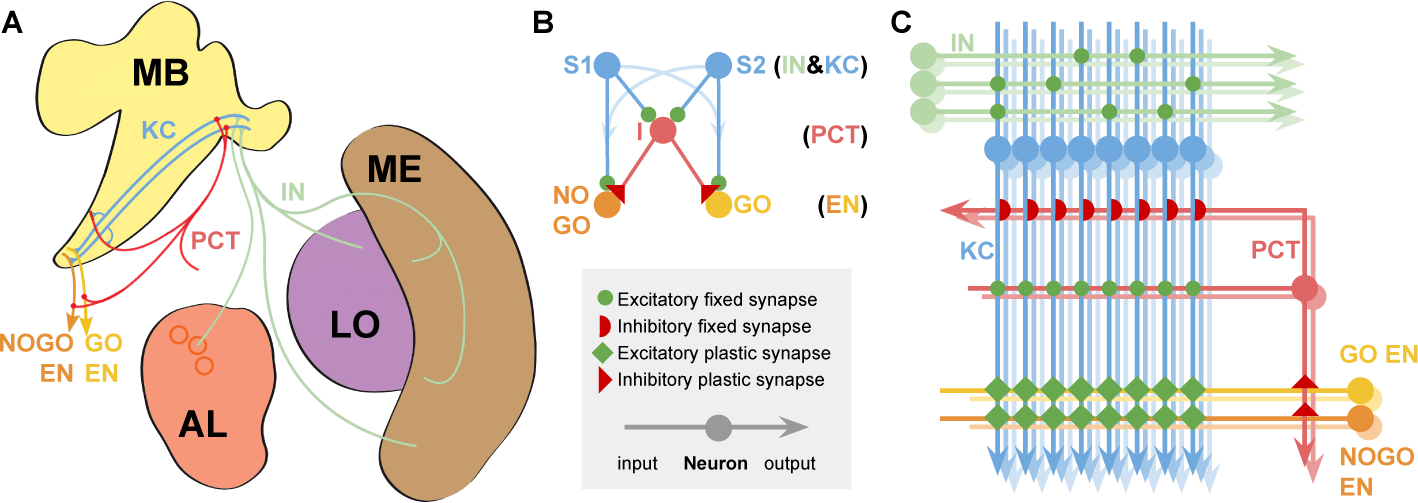
Models of the mushroom bodies based on known neuroanatomy. **A** Neuroanatomy: **MB** Mushroom Bodies; **AL** Antennal Lobe glomeruli (circles); **ME & LO** Medulla and Lobula optic neuropils. The relevant neural pathways are shown and labelled for comparison with the model. **B** Reduced model; neuron classes indicated at righthand side of sub-figure. **C** Full model, showing the model connectivity and indicating the approximate relative numbers of each neuron type. Colour coding and labels are preserved throughout all the diagrams for clarity. Excitatory and inhibitory connections indicated as in figure legend. Key of neuron types: KC, Kenyon Cells; PCT, Protocerebellar Tract neurons; IN, Input Neurons (olfactory or visual); EN, Extrinsic MB Neurons from the GO and NOGO subpopulations, where the subpopulation with the highest sum activity defines the behavioural choice in the experimental protocol (Figure 2).

The mushroom body has previously been modelled as an associative network consisting of three neural network layers (Bazhenov et al., 2013; Huerta and Nowotny, 2009), comprised of input neurons (IN) providing processed olfactory, visual and mechanosensory inputs (Mobbs, 1982; Fahrbach, 2006), an expansive middle layer of Kenyon cells (KC) which enables sparse-coding of sensory information for effective stimulus classification (Galizia, 2014), and finally mushroom body extrinsic neurons (EN) which output to premotor regions of the brain and can be considered (at this level of abstraction) to activate different possible behavioural responses (Galizia, 2014; Bazhenov et al., 2013). Here, for simplicity, we consider the EN as simply two subpopulations controlling either ‘go’ or ‘no-go’ behavioural responses only, which allow choice between different options via sequential presentations where ‘go’ chooses the currently presented option. Connections between the KC output and ENs are modifiable by synaptic plasticity (Bazhenov et al., 2013; Heisenberg, 2003; Schwaerzel et al., 2003; Strube-Bloss et al., 2011) supporting learned changes in behavioural responses to stimuli.

As outlined in the introduction, we require two computational mechanisms for solving the DMTS/DNMTS task. First is a means of storing the identity of the sample stimulus. Second is learning to use this identity to drive behaviour and solve the task. Moreover this learning must generalise to other stimuli. The computational complexity of this problem should not be underestimated; either the means of storing the identity of the sample, or the behavioural learning, must generalise to other stimulus sets. The bees were not given any reward with the transfer stimuli in Giurfa et al. (2001)’s study, so no post-training learning mechanism can explain the transfer performance. In addition, during the course of the experiment of Giurfa et al. (2001) each of the two stimuli were used as the match, i.e. for stimuli A and B the stimulus at the maze entrance were alternated between A and B throughout the training phase of the experiment. This requires, therefore, that the bees have a sense of stimulus ‘novelty’, and can associate novelty with a behaviour: either approach for DNMTS, or avoid for DMTS. With one training set the problem is solvable as delayed paired non-elemental learning tasks, however with the transfer of learning to new stimulus sets such an approach does not solve the whole task.

There is one feature of the Kenyon Cells which can fulfill this computational requirement for novelty detection, that of sensory accommodation. In honey bees, even in the absence of reward or punishment, the KC show a stark decrease in activity between initial and repeated stimulus presentations of up to 50%, an effect that persists over several minutes (Szyszka et al., 2008). This effect is also found in *Drosophila melanogaster* (Hattori et al., 2017), where there is additionally a set of mushroom body output neurons that show even starker decreases in response to repeated stimuli, and which respond to many stimuli with stimulus specific decreases (thus making them clear novelty detectors), however such a neuron has not been found in bees to date. This response decrease in Kenyon Cells found in flies and bees is sufficient to influence behaviour during a trial but, given the decay time of this effect, not likely to influence subsequent trials. The mechanism behind this accommodation property is not known, and therefore we are only able to model phenomenologically, which do by reducing the strength of the KC synapses for the sample stimulus by a fixed factor, tuned to reproduce the reduction in total KC output found by Szyszka et al. (Szyszka et al., 2008) (see Figure 3 panel E). However it should be noted that stimulus-specific adaptation is shown in many species and brain areas, and can be explained by short-term plasticity mechanisms (Tsodyks and Markram, 1997; Vasilaki and Giugliano, 2014; Esposito et al., 2014) and architectural constraints only; see for instance Yarden and Nelken (2017).

**Figure 2:**
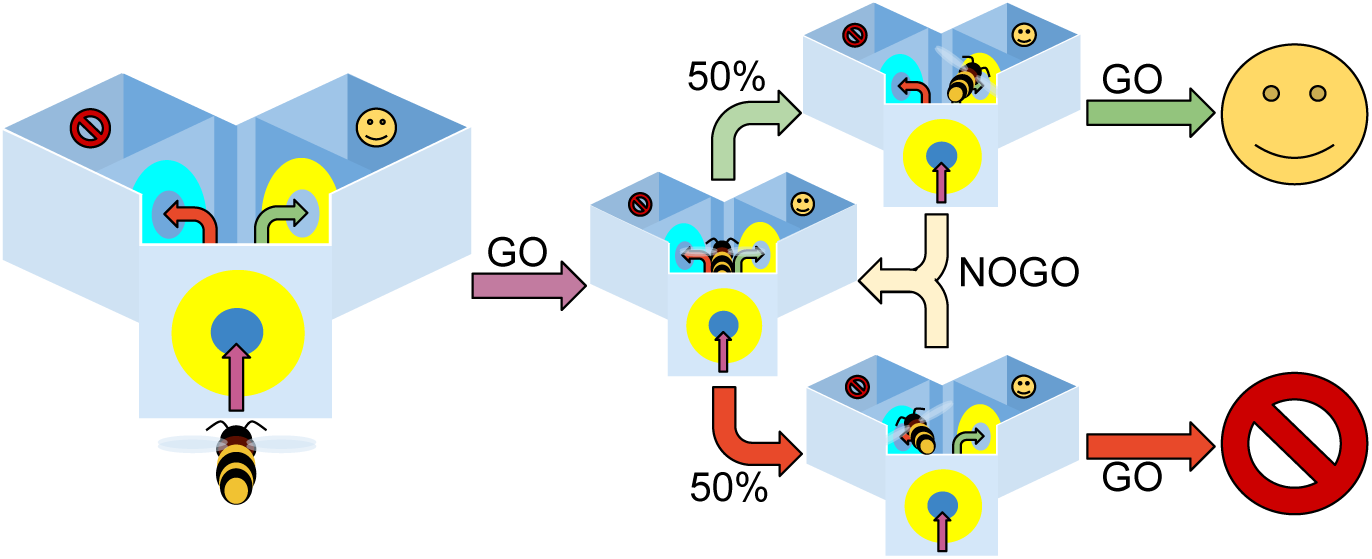
Experimental protocol for the model. The model bee is moved between a set of states which describe different locations in the Y-maze apparatus: at the entrance, in the central chamber facing the left arm, in the central chamber facing the right arm, in the left arm or in the right arm. When at the entrance or in the main chamber the bee is presented with a sensory input corresponding to one of the test stimuli; GO selection leads the bee to enter the maze when at the entrance, and to enter an arm and experience a potential reward when facing that arm; NOGO leads the bee to delay entering the maze, or to choose another maze arm uniformly at random, respectively. We can then set the test stimuli presented to match the requirements of a given trial (e.g. entrance (A), main chamber left (A), main chamber right (B) for DMTS when rewarding the left arm, or DNMTS when rewarding the right arm).

**Figure 3:**
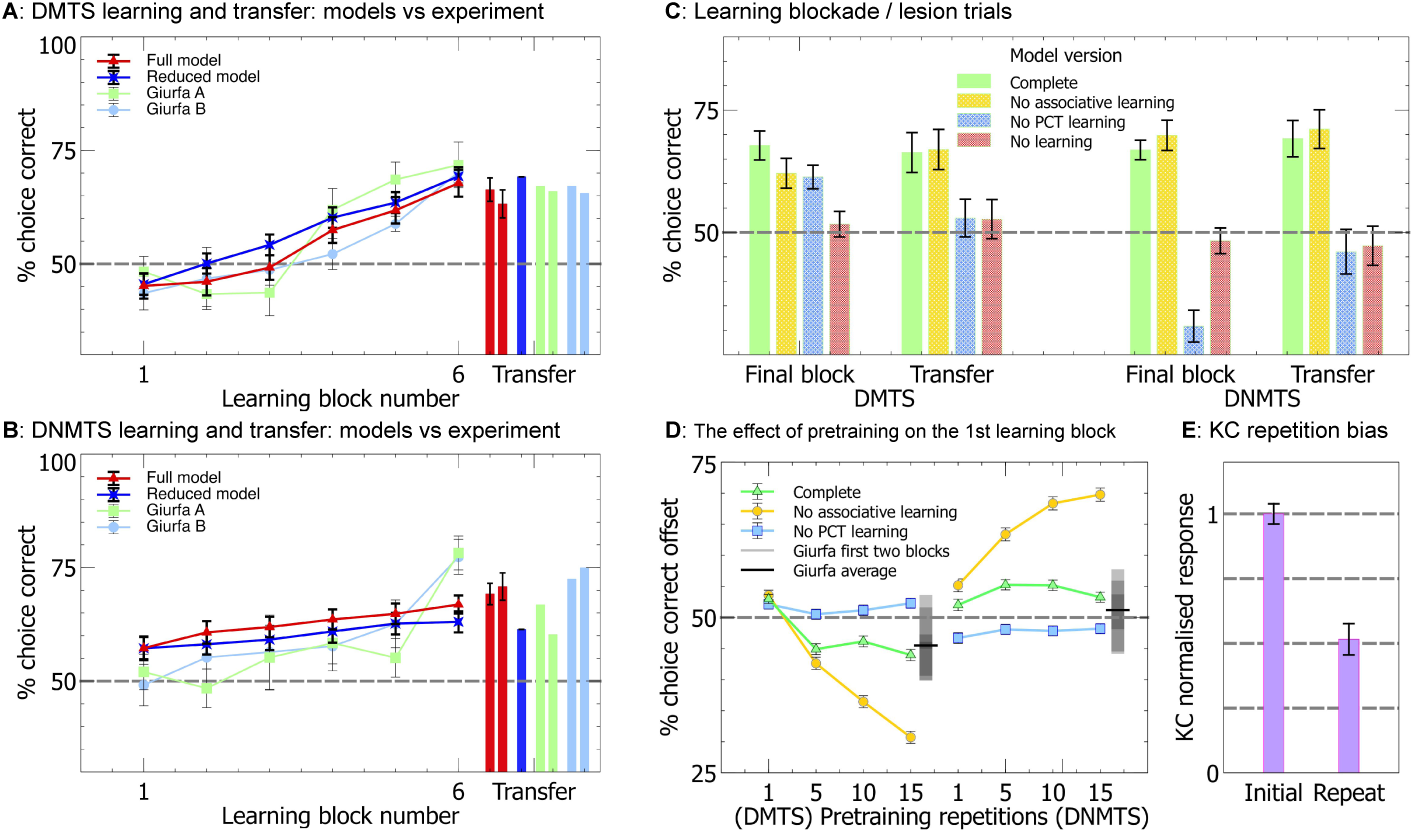
The full and reduced versions of our model reproduce the transfer of sameness and difference learning. **A & B** The average percentage of correct choices made by the model and real bees within blocks of ten trials as the task is learned (lines), along with the transfer of learning onto novel stimulus sets (bars). Both versions of the model reproduce the pattern of learning acquisition for DMTS (Full: N=338, Reduced: N= 360) and DN-MTS (Full & Reduced: N=360) found when testing real bees (test for learning: P¡0.0001), along with the transfer of learning (P¡0.0001). For DMTS *Giurfa A & B* are the data from Experiments 1 & 2 respectively from Giurfa et al. Giurfa et al. (2001), and for DNMTS *Giurfa A & B* are the data from Experiments 5 & 6 respectively from the same source. For an explanation of the initial offsets from chance for the model please see the text for panel D. **C** The blockade of plasticity from the MB and PCT pathways shows that the PCT pathway is necessary and sufficient for sameness and difference learning in the full model. All non-overlapping SEM error bars are significantly different. **D** PCT pathway learning in the absence of associative learning leads to preference for non-matching stimuli following pre-training, demonstrating that learning in the associative pathway changes the form of the sameness and difference acquisition curves. The equivalent offsets and error ranges for the first two blocks of Giurfa Experiments 1, 2, 5 & 6 along with the averages for DMTS and DN-MTS for these blocks are shown alongside the model data for comparison as overlapping grey boxes-overlapping boxes create darker regions, thus the area of greatest darkness is the point where the most of the error ranges overlap. **E** The average activity of the model KC neurons when presented with repeated stimuli.

Having identified our first computational mechanism, a memory trace in the form of reduced KC output for the repeated stimulus, we need only to identify the second, a learning mechanism that can use this reduced KC output to drive behaviour to choose the correct (matching or non-matching) arm of the y-maze. If this learning mechanism exists at the output synapses of the KC it is either specific to the stimulus - if using a pre-postsynaptic learning rule - and therefore cannot transfer, or it utilises a postsynaptic-only learning rule. Initially the postsynaptic learning rule appears a plausible solution, however we must consider that bees can learn both DMTS and DNMTS, and that learning can only occur when the bee chooses to ‘go’. This creates a contradiction, as postsynaptic learning will proportionally raise both the weaker (repeated) stimulus activity, as well as the stronger (non-repeated) stimulus activity in the GO EN subpopulation. To select ‘go’ the GO activity for the currently presented stimulus must be larger than the activity in the NOGO subpopulation, which is fixed. Therefore we face the contradiction that in the DMTS case the weaker stimulus response must be higher than the stronger one in the GO subpopulation with respect to their NOGO subpopulation counterpart responses, yet in the DNMTS case the converse must apply. No single postsynaptic learning rule can fulfil this requirement.

A separate set of neurons that can act as a relay between the KC and behaviour is therefore required to solve both DMTS and DNMTS tasks. A plausible candidate is the inhibitory neurons that form the protocerebellar tract (PCT). These neurons have been implicated in both non-elemental olfactory learning (Devaud et al., 2015) and regulatory processes at the KC input regions. They also project to the KC output regions (Ganeshina and Menzel, 2001; Haehnel and Menzel, 2010; Rybak and Menzel, 1993; Okada et al., 2007), where there are reward-linked neuromodulators and learning-related changes (Perry and Barron, 2013; Søvik et al., 2015). These neurons are few in number in comparison to the KC population, and some take input from large numbers of KC (Papadopoulou et al., 2011). We therefore propose that, in addition to their posited role in modulating and regulating the input to the KC based on overall KC activity (Papadopoulou et al., 2011), these neurons could also regulate and modulate the activity of the EN populations at the KC output regions. Such a role would allow, via synaptic plasticity, a single summation of activity from the KCs to differentially affect both their inputs and outputs. If we assume a high threshold for activity for the PCT neurons (again consistent with their proposed role) such that repeated stimuli would not activate the PCT neurons but non-repeated stimuli would, it is then possible for synaptic plasticity from the PCT neurons to the EN to solve the DMTS and DNMTS tasks and, vitally, transfer that learning to novel stimuli. We do not propose that this is the purpose of these neurons, but instead that it is a consequence of their regulatory role.

We present two models inspired by the anatomy and properties of the honey bee brain that are computationally capable of learning in DMTS and DNMTS tasks, and the generalisation of this learning to novel stimuli (Figure 1).

Our first, reduced, model is a simple demonstration that the key principles outlined above can solve DMTS and DNMTS tasks, and generalise the learning to novel stimulus sets. By simplifying the model in this way the computational principles are readily apparent. Such a simple model, however, cannot demonstrate that associative learning in the KC to EN synapses does not interfere with learning in the PCT to EN synapses or vice versa. For this we present a full model that includes the associative learning pathway from the KC to the EN, and demonstrate that this model can not only solve DMTS and DNMTS with transfer to novel stimuli, but can also solve a suite of associative learning tasks in which the MB have been implicated. The results of computational experiments performed with these models are presented below. The full model addresses the interaction of the PCT to EN learning and the KC to EN learning, as well as suggesting a possible computational role of the PCT to EN synaptic pathway in regulating the behavioural choices driven by the MB output, which we present in the Discussion.

### A reduced model of the core computational principles produces sameness and difference learning, and transfers this learning to novel stimuli

The reduced model is shown in Figure 1 Panel B and model equations are presented in Methods. The input nodes S_1_ and S_2_ represent the two alternative stimuli, where we have reduced the sparse KC representation into two non-overlapping single nodes for simplicity, and as such we do not need to model the IN input neurons separately. Node I (which corresponds to the PCT neurons, again reduced to a single node for simplicity) represents the inhibitory input to the output neurons GO and NOGO. Nodes S_1_ and S_2_ project to nodes I and to GO and NOGO with fixed excitatory weighted connection. Finally, node I projects to GO and NOGO with plastic inhibitory weighted connections. Node I is thresholded so that it only responds to novel stimuli.

Figure 3 panels A and B show the performance of the reduced model bees for task learning and transfer to novel input stimuli. While the reduced model solves the transfer of sameness and difference learning the pretraining process strongly biases the model towards non-repeated stimuli, proportional to the number of pretraining trials. Notably, this bias in the reduced model is different to that found in the full model, which we discuss below.

The model operates by adjusting the weights between the I and the GO to change the likelihood of choosing the non-matching stimulus. Since only connections from the I (representing the PCT neuron) to GO neurons are changed, the I to GO/NOGO weights are initialised to half the maximum weight value. Note that the I node is only active for the non-repeated stimulus, and this pathway has no effect for repeated stimuli. This means that if the weights are increased then non-repeated stimuli will have greater inhibition to the GO neuron, and therefore be less likely to be chosen. If the weights are decreased then non-repeated stimuli will have less inhibition to the GO neuron and therefore will be more likely to be chosen. As the conditions for changing the weights are only met when the non-repeated stimulus is chosen for ‘go‘, this means that the model only learns on unsuccessful trial for DMTS (increasing the weight), or successful trials for DNMTS (decreasing the weight).

### A full model is capable of sameness and difference learning, and transfers this learning to novel stimuli

The full model is shown in Figure 1 Panel C and model equations are presented in Methods. Figure 3 panel D shows the performance of the full model for the first block of learning following from different numbers of pretraining repetitions. When only the PCT pathway is plastic there is a large bias towards the non-repeated stimulus due to the pretraining, as found in the reduced model. This bias is reduced by the presence of the associative learning pathway, and the bias is independent of the number of pretraining trials for more than 5 trials. It should be noted that the experimental data (Giurfa et al., 2001) show indications of such a bias, in line with the results from the full model. The reduced model therefore requires fewer pretraining trials than the full model to produce a similar bias, which leads to the reduced model having large maladaptive behavioural biases for non-repeated stimuli if all stimuli are rewarded. This is important, as it suggests a role for the PCT pathway in modulating the behavioural choice of the bee. This possible role is explored further in the Discussion.

Figure 3 panels A and B show the performance of the model bees compared with the performance of real bees from Giurfa et al. (2001). In both cases the trends found in the performance of the model bees match the trends found in the real bees for both task learning and transfer to novel stimuli. It is important to note the different forms of the learning in the DMTS and DNMTS paradigms, with DNMTS slower to learn. This is a direct consequence of the inhibitory nature of the PCT neurons; excitatory neurons performing the role of the PCT neurons in the model would lead to a reversal of this feature, with DMTS learning more slowly.

### Learning in the PCT pathway of the full model is essential for transfer of learning to novel stimuli

We next sought to confirm that learning in the PCT neuron to EN pathway enabled generalisable learning of sameness and difference. Computational modelling provides powerful tools with which to do this, by comparing model performance when different elements are suppressed with the full model. We selectively suppressed the KC associative learning pathway, the PCT pathway learning, and all learning in the model. When a learning mechanism is suppressed this means that the synaptic weights stay the same throughout the training, but the pathway is otherwise active.

The results are summarised in Figure 3 panel C. It can clearly be seen that within our model learning in the PCT pathway is necessary for transfer of the sameness and difference learning to novel stimuli. Associative learning via the KC pathway alone has no effect on the transfer task performance compared to the fully learning-suppressed model. Unsuppressed associative learning leads to a preference for the matched stimulus, which has weaker KC activity, but this learning is specific to the trained stimuli, and does not transfer to novel stimuli.

### Validation: the full model is capable of performing a range of conditioning tasks

Many models have reproduced the input neuron to Kenyon Cell to Extrinsic neuron pathway (Huerta et al., 2004; Huerta and Nowotny, 2009; Bazhenov et al., 2013; Peng and Chittka, 2016), and these models demonstrate many forms of elementary and complex associative learning that have been attributed to the mushroom bodies. It is therefore important to demonstrate that in our model then PCT neuron pathway does not affect the reproduction of such learning behaviours. We therefore tested elemental and non-elemental associative learning undertaken by conditioning the PER in restrained bees, and reversal learning in free flying bees, as described in Methods. Our model is capable of reproducing the results found in experiments involving real bees, with the model’s acquisition curves showing similar to the performance to the real bees. The results are shown in Figure 4.

**Figure 4:**
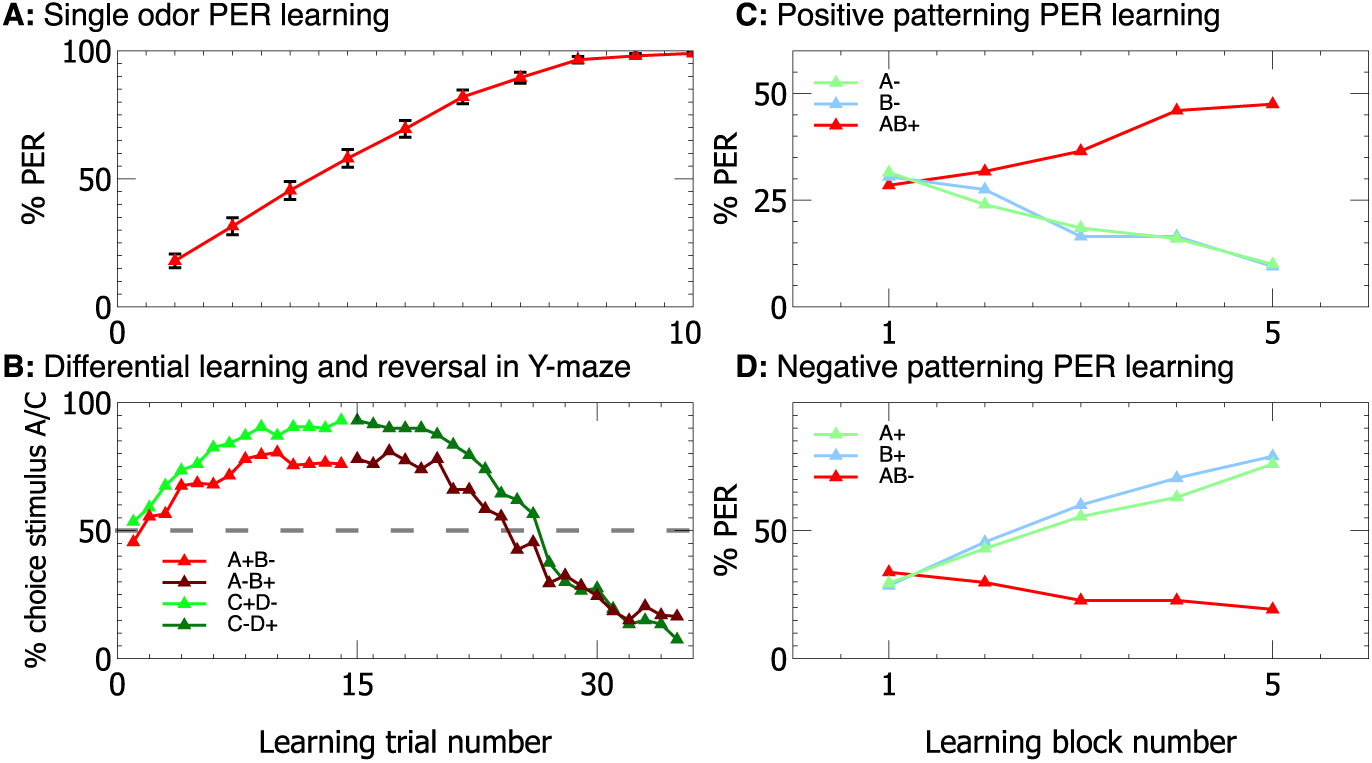
The full model is capable of performing a range of conditioning tasks. With modification of only the experimental protocol, our full model can successfully perform a range of conditioning tasks which can be performed by restrained (using the Proboscis Extention Reflex (PER) paradigm) and free flying bees. Performance closely matches experimental data with real bees (e.g. **A**:Bitterman et al. (1983), **B**:Giurfa (2004), **C & D**:Deisig et al. (2001)).

## Discussion

We have presented a simple neural model that is capable of learning the concepts of sameness and difference in Delayed Match to Sample (DMTS) and Delayed Not Match to Sample (DNMTS) tasks. Our model is inspired by the known neurobiology of the honey bee, and is capable of reproducing the performance of honey bees in a simulation of DMTS and DNMTS tasks. Our model therefore proposes a hypothesis for how animals like the honey bee might be able apparently to learn abstract concepts.

Abstract concept learning is typically described as a higher-order cognitive capacity (Wright and Katz, 2007; Avarguès-Weber and Giurfa, 2013), and one that is dependent on a top-down modulation of simpler learning mechanisms by a higher cognitive process (Moore et al., 2012). By contrast our model proposes a solution to sameness and difference learning in DMTS-style tasks with no top-down structure. The actions of the PCT neurons are integrated with the KC learning pathway and provide a parallel processing pathway sensitive to stimulus magnitude, rather than a top-down imposition of a learned concept of sameness or difference (Figure 1). This is a radical new proposal for how an abstract concept might be learned by an animal brain.

The first question we must ask when constructing a model regards plausibility. Our model (Figure 1) shows a close match to the neuroanatomical data for the mushroom bodies. Several computational requirements of our model match with experimental data, notably the sensory accommodation in the response of the KC neurons. Previous neural models based on this structure have proposed mechanisms for various forms of associative learning, including extinction of learning, and positive and negative patterning (Bazhenov et al., 2013; Arena et al., 2013; Peng and Chittka, 2016). Our model is also capable of solving a range of stimulus-specific learning tasks, including patterning (Figure 4). No plausible previous model of the MB or the insect brain has been capable of learning abstract concepts, however.

As mentioned in the Introduction, a previous model by Arena et al. (2013) demonstrates DMTS and DNMTS with transfer. Their motivation is the creation of a model for robotic implementation, rather than reproduction of behavioural observations from honey bees. While we suggest a role for the PCT neurons given experimental evidence of changes in the response of Kenyon Cells to repeasted stimuli, Arena et al.’s model assumes resonance between brain regions that is dependent upon the time after stimulus onset and the addition of specific neurons for ‘Match’ and ‘Non-match’; there is no biological evidence for either of these assumptions. Furthermore, the outcome of these additions is an increase in Kenyon cell firing in response to repeated stimuli; this is in opposition to neurophysiological evidence from multiple insect species, including honey bees (Szyszka et al., 2008; Hattori et al., 2017). In addition, Arena et al‘s proposed mechanism does not replicate the difficulty honey bees have in learning DMTS/DNMTS tasks, exhiibting learning in three trials, as opposed to 60 in real bees. In contrast, our model captures the rate and form of the learning found in real honey bees.

To enable a capacity for learning the stimulus-independent abstract concept of sameness or difference our model uniquely includes two interacting pathways. The KC pathway of the mushroom bodies retains stimulus-specific information and supports stimulus-dependent learning. The PCT pathway responds to summed activity across the KC population and is therefore largely independent of any stimulus-specific information. This allows information on stimulus magnitude, independent of stimulus specifics, to influence learning. Including a sensory accommodation property to the KCs (Szyszka et al., 2008) makes summed activity in the KCs in response to a stimulus sensitive to repetition, and therefore stimuli encountered successively (same) cause a different magnitude of KC response to novel stimuli (different) irrespective of stimulus specifics. This model is capable of learning sameness and difference rules in a simulation of the Y-maze DMTS and DNMTS tasks applied to honey bees (Figure 3), but in theory it could also solve other abstract concepts related to stimulus magnitude such as quantitative comparisons (Avarguès-Weber and Giurfa, 2013; Avarguès-Weber et al., 2014).

Our model demonstrates a bias towards non-repeated stimuli, induced by the combination of sensory accommodation in the KC neurons and PCT learning during the pretraining phase, and largely mitigated by associative learning in the KC to EN synapses. This bias (see Figure 3) is indicated in the data from Giurfa et al. (2001), and could be confirmed by further experimentation.

We note, however, that our model only supports a rather limited form of concept learning of sameness and difference. Learning in the model is dependent on sensory accommodation of the KCs to repeated stimuli (Szyszka et al., 2008). This effect is transient, and hence the capacity to learn sameness or difference will be limited to situations with a relatively short delay between sample and matching stimuli. This limitation holds for honey bee learning of DMTS tasks (Zhang et al., 2005), but many higher vertebrates do not have this limitation Lind et al. (2015). For example, in capuchins learning of sameness and difference is independent of time between sample and match (Wright and Katz, 2006). We would expect that for animals with larger brains and a developed neocortex (or equivalent) many other neural mechanisms are likely to be at play to reinforce and enhance concept learning, enabling performance that exceeds that demonstrated for honey bees. Monkey pre-frontal cortex (PFC) neurons demonstrate considerable stimulus-specificity in matching tasks, and different regions appear to have different roles in coding the salience of these stimuli (Seger and Miller, 2010; Tsujimoto et al., 2011). Recurrent neural activity between these selective PFC neurons and lower-order neural mechanisms could support such time independence. Language-trained primates did particularly well on complex identity matching tasks and the ability to form a language-related mental representation of a concept might be the reason (Premack, 1978; Premack and Premack, 1983; Thompson and Oden, 1995).

Wright and Katz (Wright and Katz, 2007) have utilised a more elaborate form of a MTS task in which vertebrates simultaneously learn to respond to sameness and difference, and are trained with large sets of stimuli rather than just two. They argue this gives less-ambiguous evidence of true concept learning since both sameness and difference are learned during training, and the large size of the training stimulus set encourages true generalisation of the concept. In theory our model could also solve this form of task, but it is unlikely a honey bee could. Capuchins, rhesus and pigeons required hundreds of learning trials to learn and generalise the sameness and difference concepts (Wright and Katz, 2007). Bees would not live long enough to complete this training,

Finally as a consequence of our model we question whether it is necessary to consider abstract concept learning to be a higher cognitive process. Mechanisms necessary to support it may not be much more complex than those needed for simple associative learning. This is important because many behavioural scientists still adhere to something like Lloyd Morgan’s Canon (Lloyd Morgan, 1903), which proposes that “in no case is an animal activity to be interpreted in terms of higher psychological processes if it can be fairly interpreted in terms of processes which stand lower in the scale of psychological evolution and development” (Lloyd Morgan (1903) p59). Yet the Canon is therefore reliant on an unambiguous stratification of cognitive processes according to evolutionary history and development (Sober, 2015). If abstract concept learning is in fact developmentally quite simple, evolutionarily old and phylogenetically widespread, then Morgan’s Canon would simply beg the question of why even more animals do not have this capacity (Mikhalevich, 2015). We argue that far more information on the precise neural mechanisms of different cognitive processes, and the distribution of cognitive abilities across animal groups, is needed in order to properly rank capacities as higher or lower.

## Methods

### Model parameter selection

Many of the parameters of the model were fixed by the neuroanatomy of the honey bee, as well as the previous values and procedures described in Bazhenov et al. (2013), with the following modifications.

First, we increased the sparseness of the connectivity from the PN to the KC.

Second, the reduction in the magnitude of the KC output to repeated stimuli was tuned to replicate the magnitude of reduction described in Szyszka et al. (2008).

Third, the learning rates were set so that acquisition of a single stimulus is rapid. In addition there are two ratios from this initial value that must be set. These are the ratio of the speed of excitatory associative learning in the Kenyon Cell to Extrinsic Neuron pathway to the inhibitory learning in the Protocerebellar Tract to Extrinsic Neuron pathway, and the ratio of the speed of acquisition when rewarded to the speed to extinction when no reward is given. We conservatively set both of these ratios to 2:1, with excitatory learning faster than inhibitory learning, and extinction faster than acquisition.

Finally, we tuned the threshold value for the PCT neurons so that they only responded to a new stimulus, and not a repeated one.

A full list of the parameters can be found in Table 1.

**Table 1:** Model parameters; all parameters are in arbitrary units

| Name | Value | Name | Value |
| --- | --- | --- | --- |
| **Full** |  |  |  |
| $N_{IN}$ | 144 | $N_{KC}$ | 5000 |
| $N_{EN}$ | 8 | $N_{PCT}$ | 6 |
| $p_{IN \rightarrow KC}$ | 0.02 | | |
| $b$ | 1.2 | $b_s$ ($l > 0$) | 150 |
| $b_s$ ($l = 0$) | 120 | | |
| **Reduced** |  |  |  |
| $c$ | 80 | $d_0$ | 1 |
| **Shared** |  |  |  |
| $\lambda_e$ | 0.06 | $\lambda_i$ | 0.03 |
| $R_b$ | 2/3 | | |

### Reduced Model

The reduced model is shown in Figure 1, and described in the text in Results. Here are the equations governing the model.

The input node S_1_ projects to node I via a fixed excitatory weight of 1.0 and to GO and NOGO with excitatory weights 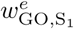 and 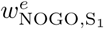 correspondingly (superscript denotes *excitatory* and subscript the connection from neuron *S*_1_ to G*O/N OGO*). Similarly, node S_2_ projects to I via an excitatory weight 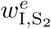 and to GO and NOGO with excitatory weights 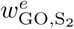 and 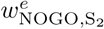. Finally, node I projects to GO and NOGO with inhibitory weights 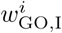 and 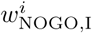 correspondingly. Node I is a threshold linear neuron with a cut-off at high values of activity *x*_*max*_. Nodes GO and NOGO are linear neurons, with activities restricted between

The model is described by the following equations, where only one input node S_1_ or S_2_ are active (but not both, as the bee observes one option at a time), where the activities of neurons I, GO and NOGO are calculated by:

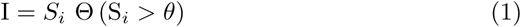

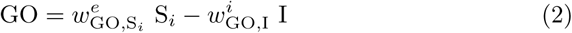

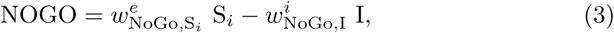

where *i* = 1, 2 is an index taking values depending on which stimulus is present (S_1_ or S_2_), and neuronal activities of I, GO and NOGO are constrained between *x*_*min*_ and *x*_*max*_. If a stimulus has been shown twice, during its second presentation there is a suppression of the neuronal activity that represents the specific stimulus, consistent with experimental findings Szyszka et al. (2008). This is modelled as a reduction by a factor of 0.7 of the value *S*_*i*_ for the repeated stimuli.

To calculate the proabability of the behavioral outcome of GO or NOGO being the winner we use the following equation:

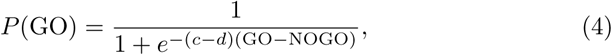

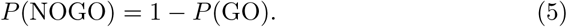

where *c* is a fixed coefficient and *d* a bias that increases linearly with the time it takes to make a decision, in the following way:

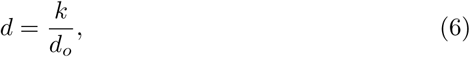

with *k* being the number of consequent iterations for which GO has not been selected, set at zero at the beginning of each stimulus presentation. The parameter *d*_*o*_ is a constant, and selected so that the factor *c – d* will always be positive. This parameterisation makes sure that the longer it takes for a decision GO to be made, the higher the probability that GO will be chosen at the next step.

Inhibitory synaptic weights *w*^*i*^ are learned using the following equation:

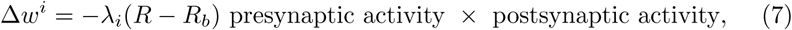

where *λ*_*i*_ is the learning rate of the inhibitory weights, reward *R* = 1 if reward is given, and zero in all other cases, *R*_*b*_ is a reward baseline, and the presynaptic (postsynaptic) activity is 1 if the presynaptic (postsynaptic) neuron is active and 0 elsewhere. This is a reward-modulated Hebbian rule also known as a three factor rule Vasilaki et al. (2009); Richmond et al. (2011).

Additional neuronal inputs with similar connectivity as *S*_1_ and *S*_2_, not shown explicitly in the diagram, are also present in the model simulations, and the constructing the equations for these simply requires substitution of *S*_*i*_ for *T*_*i*_ in Equations 1, 2 and 3. These represent the transfer stimuli and can be used following training to demonstrate transfer of learning. Details of training the model can be found in the Experiment subsection of the Methods.

### Full Model

The full model is shown in Figure 1. Our model builds on a well established abstraction of the mushroom body circuit (see Huerta et al. (2004); Huerta and Nowotny (2009); Bazhenov et al. (2013)) to model simple learning tasks.

The main structure of the model consists of an associative network with three neural network layers. Adapting terminology and features from the insect brain we label these: input neurons (IN) (correponding to S in the Reduced Model), a large middle layer of Kenyon cells (KC) (correponding to the S to GO / NOGO connections in the Reduced Model), and a small output population of mushroom body extrinsic neurons (EN) separated into GO and NOGO subsets (as in the Reduced Model). The connections, *c*_*ij*_, between the IN and KC are fixed, and are randomly selected from the complete connection matrix with a fixed probability *p*_*IN−>KC*_ = 0.02. Connections from the KC to the EN are plastic, consisting of a fully connected matrix. The connection strength between the *j*th KC and the *k*th EN (*w*_*jk*_) can take a value between zero and one. The neural description used in the entire model is linear with a bottom threshold, and contains no dynamics, consisting of a summation over the inputs followed by thresholding at a value *b* via a Heaviside function Θ, with a linear response above the threshold value. This gives the associative model as the following, where the outputs of *i*th, *j*th and *k*th neurons of the IN, KC and EN populations are *x*_*i*_, *y*_*j*_ and *z*_*k*_ respectively, and *M* describes the modulation of KC activity for the stimulus seen at the maze entrance:

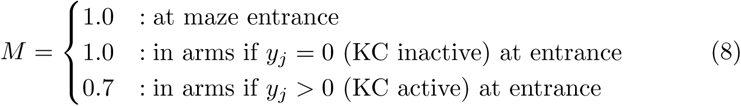

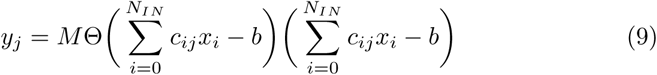

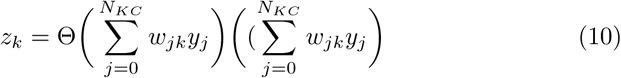

where *N*_*IN*_ is the number of IN and *N*_*KC*_ is the number of KC.

DMTS generalisation is performed by the inhibitory protocerebral tract (PCT) neurons *s*_*l*_ (correponding to I in the Reduced Model) described by the following equations:

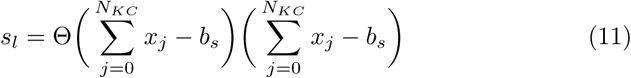

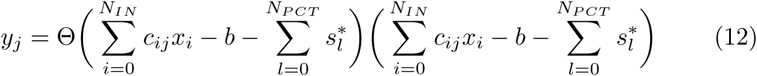

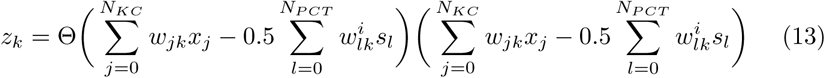

Where *w*_*lk*_ are the inhibitory weights between the *P CT* neurons. The *\** denotes 10 iteration delayed activity from the PCT neurons due to delays in the KC-*>*PCT-*>*KC loop.

Learning takes place according to equation (7), and the following equation for excitatory synapses:

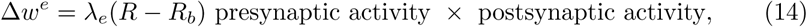

where *λ*_*e*_ is the learning rate of the excitatory weights, reward *R* = 1 if reward is given, and zero in all other cases, *R*_*b*_ is a reward baseline, and the presynaptic (postsynaptic) activity is 1 if the presynaptic (postsynaptic) neuron is active and 0 elsewhere.

Similarly to the reduced model, a decision is made when the GO EN subpopulation activity is greater than the NOGO EN population by a bias *Rd*, where *d* increases every time a NOGO decision is made by 10.0, and R is a uniform random number in the range [−0.5, 0.5]. To prevent early decisions the sum of the whole EN population activity must be greater than 0.1.

### Experiment

Our challenge is to reproduce Giurfa et al’s data demonstrating bees solving DMTS and DNMTS tasks (Giurfa et al., 2001). To aid exploration of our model we simplify the task it must face, while retaining the key elements of the problem as faced by the honeybee. We therefore embody our model in a world described by a state machine. This simple world sidesteps several navigation problems associated with the real world, however we believe that for the sufficiency proof we present here such simplifications are acceptable-the ability of the honeybee to form distinct and consistent neural representations of the training set stimuli as it flies through the maze is a prerequisite of solving the task, and is therefore assumed.

The experimental paradigm for our Y-maze task is shown in Figure 2. The model bee is moved between a set of states which describe different locations in the Y-maze apparatus: at the entrance, in the central chamber facing the left arm, in the central chamber facing the right arm, in the left arm or in the right arm. When at the entrance or in the main chamber the bee is presented with a sensory input corresponding to one of the test stimuli. We can then set the test stimuli presented to match the requirements of a given trial (e.g. entrance (A), main chamber left (A), main chamber right (B) for DMTS when rewarding the left arm, or DNMTS when rewarding the right arm).

### Experimental environment

The experimental environment consists of a simplified Y-maze (see Figure 2: main paper), in which the model bee can assume one of three positions: at the entrance to the Y-maze; at the choice point in front of the left arm; at the choice point in front of the right arm. At each position there are two choices available to the model: go and no-go. Go is always chosen at the entrance to the Y-maze as bees that refuse to enter the maze would not continue the experiment. Following this there is a random choice of one of the two maze arms, left or right. If the model chooses no-go this procedure is repeated until the model chooses to go. As no learning occurs at this stage it is possible for the model to constantly move between the two arms, never choosing to go. To prevent this eventuality we introduce a Uniformly distributed random bias *B* to the go channel that increases with the number of times the model chooses no-go 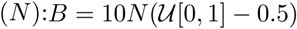.

The IN neurons are divided into non-overlapping groups of 8 neurons, each representing a stimulus. These are:

- Z: Stimulus for pretraining
- A: Stimulus for training pair
- B: Stimulus for training pair
- C: Stimulus for transfer test pair
- D: Stimulus for transfer test pair
- E: Stimulus for second transfer test pair
- F: Stimulus for second transfer test pair

Each group contains neurons which are zero when the stimulus is not present, and a value of 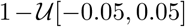 - consistent across the experiment for each bee, but not between bees - when active.

### DMTS / DNMTS experimental procedure

#### Models as animals

We use the ‘models as animals’ principle to reproduce the experimental findings of Giurfa et al. (2001), creating many variants of the model which perform differently in the experiments. To do this we change the random seed used to generate the connectivity *c*_*ij*_ between the IN and the KC neurons. For these experiments we use 360 model bee variants, each of which is individually tested, as this matches the number of bees in Giurfa et al. (2001).

#### Pretraining familiarisation

As is undertaken in the experiments with real bees, we first familiarise our naive model bees with the experimental apparatus. This is done by first training ten rewarded repetitions of the bee entering the Y-maze with a stimulus not used in the experiment. In these cases the model does not choose between go and no-go, it is assumed that the first repetition represents the model finding the Y-maze and being heavily enough rewarded to complete the remaining repetitions. Following these ten repetitions the bee is trained with ten repetitions to travel to each of the two arms of the Y-maze. This procedure ensures that the bees will enter the maze and the two arms when the training begins, allowing them to learn the task.

#### Training

The training procedure comprises 60 trials in total, divided into blocks of 10 trials. The protocol involves a repeated set of four trials: two trials with each stimulus at the maze entrance, with each of these two trials having the stimulus at the maze entrance on different arms of the apparatus. In the case of match-to-sample the entrance stimulus is rewarded and the non-entrance stimulus is punished, and vice versa for not-match-to-stimulus.

#### Transfer test

For the transfer test we do not provide a reward or punishment, and test the models using the procedure for Training, substituting the transfer test stimuli for the training stimuli. Two sets of transfer stimuli are used, and four repetitions (left and right arm with each stimulus) are used for each set of stimuli.

### Testing performance of the full model in other conditioning tasks

In addition to solving the DMTS and DNMTS tasks, we must validate that our proposed model can also perform a set of conditioning tasks that are associated with the mushroom bodies in bees, without our additional PCT circuits affecting performance. Importantly, these tasks are all performed with exactly the same model parameters that are used in the DMTS and DNMTS tasks, yet match the timescales and relative performances found in experiments performed on real bees. We choose four tasks, which comprise olfactory learning experiments using the proboscis extention reflex (PER) that are performed on restrained bees as well as visual learning experiments performed with free flying bees (Figure 4).

#### Differential learning / reversal experimental procedure (Figure 4, panel B)

These experiments follow the same protocol as the DMTS experiments, except that for the first fifteen trials one stimulus is always rewarded when the associated arm is chosen (no reward or punishment is given for choosing the non-associated arm), and subsequent to trial fifteen the other stimulus is rewarded when the associated arm is chosen. No pretraining or transfer trials are performed and the data is analysed for each trial rather than in blocks of 10 due to the speed of learning acquisition. 200 virtual bees are used for this experiment (see Figure 4, panel B for results, to be compared with Giurfa (2004)).

#### Proboscis Extension Reflex (PER) Experiments

The Proboscis Extension Reflex (PER) is a classical conditioning experimental paradigm used with restrained bees. In this paradigm the bees are immobilised in small metal tubes with only the head and antennae exposed. Olfactory stimuli (conditioned stimuli) are then presented to the restrained bees in association with a sucrose solution reward (unconditioned stimulus) (see Bitterman et al. (1983) for full details).

For the PER experiments we separate the IN neurons as described in Section, however as the bees are restrained for these experiments we present odors following a pre-defined protocol, and the choices of the bee do not affect this protocol.

#### Single odor learning experimental procedure (Figure 4, panel A)

In the single odor experiments we use the procedure outlined in Bitterman et al Bitterman et al. (1983). In this procedure acquisition and testing occur simultaneously. The real bees are presented an odor, and after a delay rewarded with sucrose solution. If the animal extends its proboscis within the delay period it is rewarded directly and considered to have responded, if it does not the PER is invoked by touching the sucrose solution to the antennae and the animal is rewarded but considered not to have responded. To match this protocol the performance of the model was recorded at each trial, with NOGO considered a failure to respond to the stimulus, and GO a response. At each trial a reward was given regardless of the model’s performance.

#### Positive / negative patterning learning experimental procedure (Figure 4, panels C & D)

In these experiments we follow the protocol described in Deisig et al. (2001). We divide the training into blocks, each containing four presentations of an odor or odor combination. For positive patterning we do not reward individual odors A and B, but reward the combination AB (A-B-AB+). In negative patterning we reward the odors A and B, but not the combination AB (A+B+AB-). In both cases the combined odor is presented twice for each presentation of the individual odors, so a block for positive patterning is [A-,AB+,B-,AB+] for example, while for negative patterning a block is [A+,AB-,B+,AB-]. Performance is assessed as for the single odor learning experiment, with the two combined odor responses averaged within each block.

### Software and implementation

The reduced model was simulated in GNU Octave (John W. Eaton David Bateman and Wehbring, 2015). The full model was created and simulated using the SpineML toolchain (Richmond et al., 2013) and the SpineCreator graphical user interface (Cope et al., 2015). These tools are open source and installation and usage information can be found on the SpineML website at http://spineml.github.io/. Input vectors for the IN neurons and the state engine for navigatation of the Y-maze apparatus are simulated using a custom script written in the Python programming language (Python Software Foundation, https://www.python.org/) interfaced to the model over a TCP/IP connection.

Statistical tests were performed as in Giurfa et al. (2001) using 2×2 *X*^2^ tests performed in *R* (R Core Team, 2013) using the chisq.test() function.

The code is available online at http://github.com/BrainsOnBoard/bee-concept-learning

## Acknowledgements

We thank Martin Giurfa, Thomas Nowotny and James Bennett for their constructive comments on the manuscript. JARM and EV acknowledge support from the Engineering and Physical Sciences Research Council (grant numbers EP/J019534/1 and EP/P006094/1). JARM and ABB acknowledge support from a Royal Society International Exchanges Grant. ABB is supported by an Australian Research Council Future Fellowship Grant 140100452 and Australian Research Council Discovery Project Grant DP150101172.

